# Community needs for FAIR pathogen data: a cross-sectional survey of infectious-disease researchers and educators

**DOI:** 10.64898/2026.04.14.718420

**Authors:** Geert van Geest, Daniel Thomas-Lopez, Anna A. Feitzinger, Lily A. Weissgold, Sam Halabi, Isabel Cuesta, Erik Hjerde, Kim Tamara Gurwitz, Nishtha Arora, Aitana Neves, Patricia M. Palagi, Jason J. Williams

**Affiliations:** SIB Swiss Institute of Bioinformatics, Lausanne, Switzerland; EMBL’s European Bioinformatics Institute, Cambridge, United Kingdom; Cold Spring Harbor Laboratory, Cold Spring Harbor, NY, USA; Leibniz Institute DSMZ, Braunschweig, Germany; Center for Transformational Health Law, Georgetown University, Georgetown, Washington D.C., USA; Bioinformatics Unit, Institute of Health Carlos III, Madrid, Spain; The Arctic University of Norway, Tromsø, Norway

## Abstract

**Background:** Datasets related to infectious diseases are essential for public health decision-making, yet their reuse remains limited by persistent barriers to data sharing and integration. Achieving data that are Findable, Accessible, Interoperable, and Reusable (FAIR) is widely recognized as essential for accelerating scientific discovery and enabling coordinated responses to emerging threats, but the needs of the global pathogen data community have not been systematically characterized.

**Aim:** This study, conducted by the Pathogen Data Network (PDN), aims to identify infrastructural and educational priorities among stakeholders working with infectious disease-related data—namely researchers, educators, healthcare professionals, bioinformaticians, and policymakers who generate, curate, or use pathogen data, in order to guide community-responsive support for data sharing and interoperability.

**Methods:** A cross-sectional stakeholder survey was disseminated as an open, non-probability (convenience) survey to prospective participants reached through PDN networks, targeted mailing lists, and open professional and social-media channels. A total of 136 responses from researchers, healthcare professionals, bioinformaticians, and educators were analyzed descriptively to identify prioritized barriers, training needs, and preferred support mechanisms.

**Results:** Respondents consistently identified structural constraints as the primary impediments to effective data use, including limited funding (74%), data-aggregation challenges (68%), and a shortage of skilled personnel (52%). Respondents identified bioinformatics for infectious disease research (68%) as the highest priority for training, followed by guidance on using the integrated pathogen data and tools portal provided by the PDN, the Pathogens Portal (51%). The Pathogens Portal was also ranked as the most essential PDN resource (72%). Preferred training formats included virtual short courses (68%) and webinars (66%). Notably, while researchers emphasized technical subjects like machine learning, educators prioritized foundational case studies.

**Conclusion:** These findings provide an evidence-based descriptive summary of community needs and suggest that barriers to FAIR pathogen data are predominantly systemic rather than purely technological. The survey framework and openly available dataset offer a reusable template for assessing needs in other communities and regions. By aligning training, infrastructure development, and outreach with empirically identified priorities, organizations supporting infectious disease research can strengthen the interoperability and reuse of data and establish a benchmark for future community-driven improvements.

## Introduction

Datasets related to infectious disease are fundamental to public health decision making, especially in the age of genomic medicine. Yet the full potential of these datasets remains unused when they are siloed, poorly annotated, or inaccessible to the broader scientific community [1,2]. Implementing the FAIR principles [3] - making data Findable, Accessible, Interoperable, and Reusable - addresses these barriers by standardizing metadata, ensuring reliable storage, and facilitating sharing and integration across disciplines and borders. When implemented effectively, FAIR data can accelerate scientific discovery and support timely, evidence-based responses to emerging health threats.

Despite general support for data sharing among researchers [4], Infectious disease datasets remain difficult to share and reuse in practice [1]. Barriers extend beyond technical constraints [2] and include insufficient institutional support, limited expertise to complete submission processes, data sensitivity and ethical considerations, and limited computational expertise [1,4,5]. Many of these barriers can be overcome with targeted education, training [5] and raising awareness. Training can be incorporated at a variety of levels and formats, ranging from degree programs to short-format training [6], and the preferred approach depends on the geographical region, required competencies [6], available resources, and domain. Recent mapping of pathogen genomics training provision shows that courses are predominantly short and introductory and often lack structured assessment [7], yet it remains unclear how well this supply matches what practitioners report needing. Because training needs and resource constraints vary across regions, disciplines, and professional roles, a systematic understanding of community priorities is essential for designing effective interventions that improve data sharing and interoperability.

The Pathogen Data Network (PDN) is an international consortium that aims to build a robust, FAIR ecosystem for infectious-disease data. In the 1^st^ year, PDN has focused on developing open-source tools, an integrated portal to connect pathogen data and tools (Pathogens Portal), targeted training, and community outreach. A central component of PDN’s strategy is to serve the needs of the diverse stakeholders who generate, curate, and rely on pathogen data. To capture these perspectives, we designed a comprehensive cross-sectional stakeholder survey aimed at individuals whose work involves data relevant to human health during PDN’s 1^st^ year (2024-2025). The survey assessed perceived barriers, resource priorities, and training needs to inform community-responsive support strategies.

The questionnaire was disseminated via email and social media to roughly 10,000 recipients. This was an open online survey using convenience sampling. Completed responses provide a cross-sectional snapshot of experiences and expectations within a specialized expert community, establishing an empirical baseline for aligning infrastructure, training, and outreach with user needs. Here, we present the survey findings and discuss their implications for strengthening FAIR-aligned stewardship and reuse of data.

## Material and Methods

### Survey Design and Objectives

A cross-sectional stakeholder survey was developed to characterize barriers that researchers, educators, and related professionals encounter when using pathogen data in research or educational contexts, and to identify community priorities for training and infrastructure. The instrument was designed collaboratively by members of the PDN working group and reviewed internally for clarity and relevance before dissemination. Questions were developed de novo based on PDN strategic goals and previous consultations with community partners rather than adapt ed from existing validated tools.

The study protocol and survey instrument were reviewed and determined exempt by the institutional review board of Cold Spring Harbor Laboratory (IRB-25-4). Participants gave informed consent; no compensation was offered.

The final questionnaire was comprised of 10 items, including multiple-choice, categorical, and open-ended questions. Items were organized into sections addressing: (1) respondent demographics and professional role, (2) institutional and geographic context, (3) perceived barriers to working with pathogen data, and (4) training and resource needs. Questions assessing barriers used a three-point scale (‘Significant barrier’, ‘Not significant barrier’, ‘Not applicable’). Reporting of this survey was guided by relevant items from the CROSS [8] checklist for survey-based research. The survey questions are available from: https://doi.org/10.5281/zenodo.19494321.

### Participant Recruitment and Data Collection

The survey was administered online through SurveyMonkey. It was announced on February 20, 2025, and remained open until April 25, 2025. Recruitment was conducted through multiple channels to maximize reach across the global infectious-disease research and education community. Two large email campaigns reached approximately 10,000 individuals, including 7,000 higher-education faculty working in pathogen-related disciplines and 4,000 active awardees of the US National Institutes of Health’s National Institute of Allergy and Infectious Diseases (NIAID). Additional invitations were circulated via the PDN mailing list, professional networks, social-media platforms, and targeted outreach to international health research and policy organizations.

To maximize representation, PDN collaborators compiled a contact list of more than 70 organizations and initiatives involved in pathogen data sharing, curation, and training. Contacts represented a mix of governmental organizations, mission-driven consortia, data repositories, academic institutions, and professional societies across multiple continents. This list included entities such as the U.S. Centers for Disease Control and Prevention (CDC), European Centre for Disease Prevention and Control (ECDC), Food and Drug Administration (FDA), World Health Organization (WHO) International Pathogen Surveillance Network, Africa CDC, European Virus Bioinformatics Center, Global Microbial Identifier, and several national public-health institutes, academic centers, and professional societies (e.g., American Society for Microbiology, Sociedade Brasileira de Microbiologia, European Society of Clinical Microbiology and Infectious Diseases). We were unable to capture the extent to which organizations may have shared the survey, though this presumably varied.

To confirm eligibility, respondents were required to affirm that their work or study involves pathogen data relevant to human health. No personal identifying information was collected.

### Data Analysis

Quantitative responses were analyzed descriptively to summarize response frequencies and proportions. Only responses that answered one or more questions past question 3 (‘Which of the following best describes your primary role in working with pathogen data?’) were marked as ‘valid’ and considered for downstream analysis. The analysis assumes missing data are completely at random (MCAR). The goal of analysis was to characterize relative frequencies and majority responses rather than to test statistical hypotheses. Open-ended responses were reviewed qualitatively to identify recurring themes. The analysis was performed using R version 4.5.1 [9]. The visualizations were generated with the R packages ggplot [10], cowplot [11] and rnaturalearth [12]. The analysis code is available from: https://github.com/sib-swiss/pdn_survey_analysis and anonymized raw data from: https://doi.org/10.5281/zenodo.19494321.

## Results

A total of 198 participants initiated the Global Survey of Pathogen Data Practices between March and April 2025 of which 136 individuals were considered ‘valid’ (responded to questions past the demographic questions that were at the beginning of the survey). A precise response rate could not be calculated; the estimate of ∼10,000 recipients reflects only the direct email campaigns described in the Methods section. Respondents represented a broad spectrum of professional roles (Fig 1A). The majority identified themselves as researchers, analysts, or health-care professionals (58 %; n = 79), followed by educators (26 %; n = 36), developers (7 %; n = 9), policymakers (4 %; n = 5) and data stewards (3 %; n = 4). Geographically (Fig 1B), participants were concentrated in the United States (57 %; n = 72) with additional representation from Switzerland (7 %; n = 9), Norway (6 %; n = 8), the United Kingdom (6 %; n = 7) and India (3 %; n = 4); the remaining 21 % were spread across other countries.

**Fig 1.**
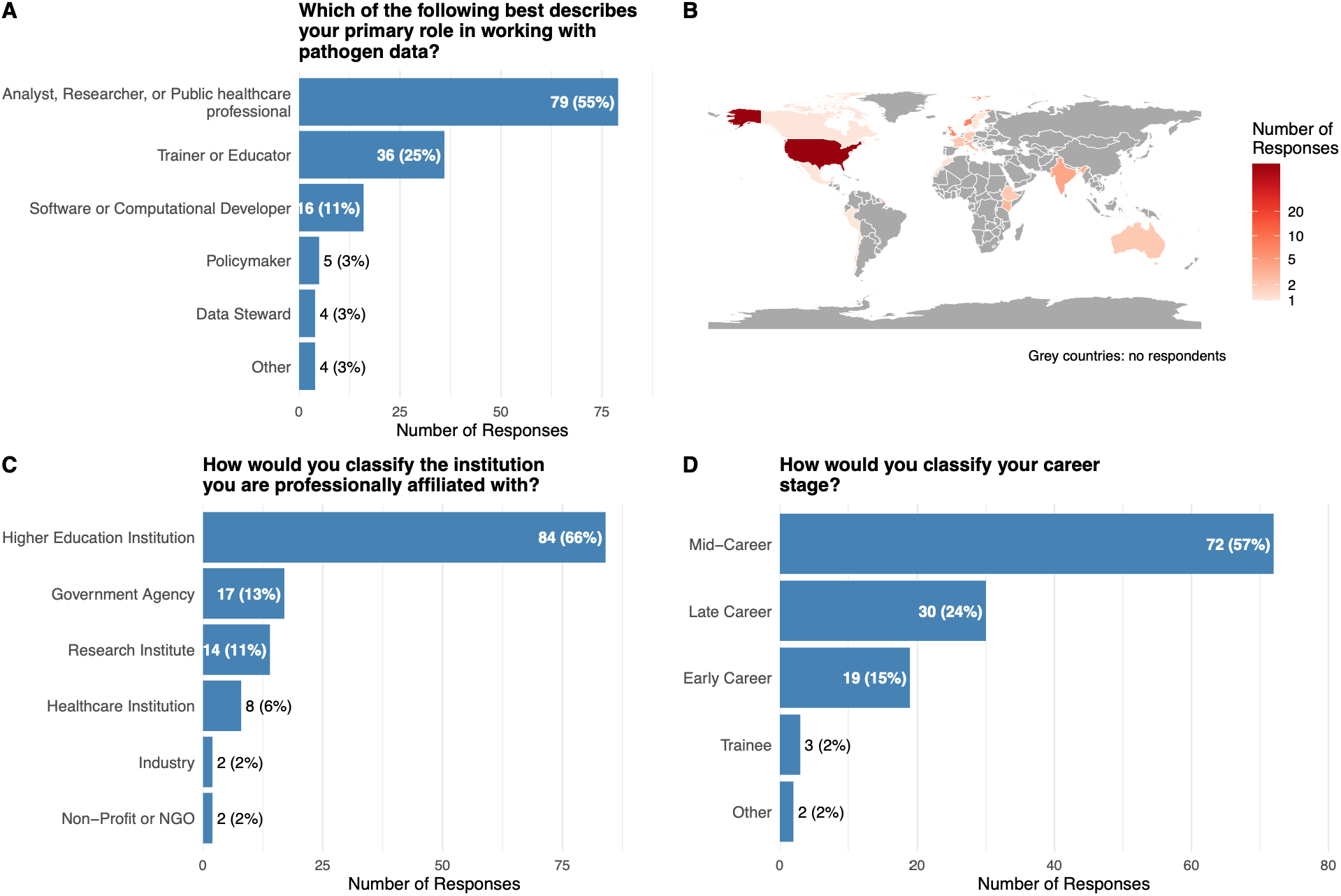
Respondent characteristics: the primary role (A), the country of work (B), type of institution (C) and career stage (D).

Most respondents were affiliated (Fig 1C) with higher-education institutions (66 %; n = 84); the remainder worked for government agencies (13 %; n = 17), research institutes (11 %; n = 14), industry (2 %; n = 2) or non-profit organizations (2 %; n = 2). In terms of career stage (Fig 1D): mid-career professionals formed the largest group (57 %; n = 72), followed by late-career respondents (24 %; n = 30), early-career professionals (15 %; n = 19), trainees (2 %; n = 3) and retired or “other” respondents (2 %; n = 2).

When asked to evaluate barriers to working effectively with pathogen data (Fig 2A), three issues emerged as the most significant. Limited funding was the most frequently cited barrier, with 74 % of participants indicating that insufficient budgets for experiments, computational tools and subscription-based resources hampered their work. Data-aggregation challenges, -specifically the difficulty of combining and sharing data across siloed systems and organizations, -were identified by 68 % of respondents, while a shortage of skilled personnel affected 52 % of the sample. Conversely, most respondents regarded several potential obstacles as not significant. Inadequate recognition for data contributions was deemed unimportant by 62 %, insufficient technical resources by 61 % and the potential for data misuse by 60 %. When participants were asked to name the single most important barrier, restricted access to data tied with limited funding; each of these two factors was selected by 16 % of respondents (Fig 2B).

**Fig 2.**
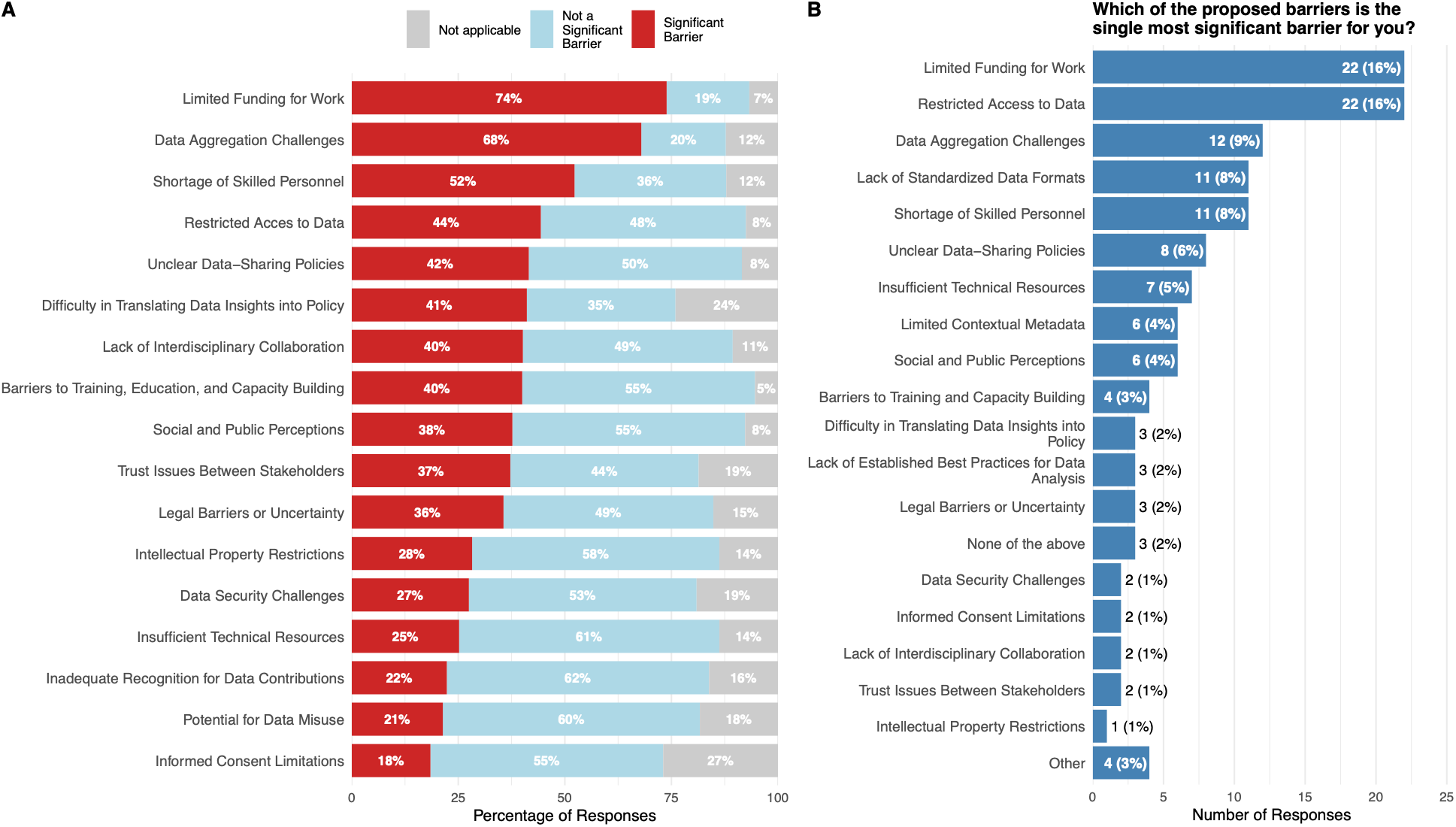
Overview of barriers to pathogen data sharing. Answer to the question ‘What are the barriers you face when working with pathogen data’ by each of the categories (A) and the single most important category (B).

Training needs were also explored (Fig 3). The most frequently selected priority area was bioinformatics for infectious-disease research, which 68 % of respondents marked as a high-priority topic. Guidance on the use of the Pathogens Portal, and training in visualization and communication of results were each chosen by roughly half of the participants (51 % and 50 %, respectively). Additional topics of interest included infectious-disease data management (41 %) and the application of machine-learning and artificial-intelligence methods to pathogen research (39 %). A direct comparison of researchers (n = 106) and educators (n = 49) revealed clear differences in emphasis. When the responses were examined by professional role (Fig 3B), researchers placed particular emphasis on technical and applied subjects:47 % expressed interest in machine-learning applications, 32 % indicated data-management planning as a priority, 29 % highlighted FAIR-data principles and 20 % selected international standards or regulations. Educators, by contrast, favored more foundational content: 71 % selected real-world case-studies, 41 % valued pathogen-domain knowledge and conceptual overviews, and none (0 %) chose international standards or regulations as a priority.

**Fig 3.**
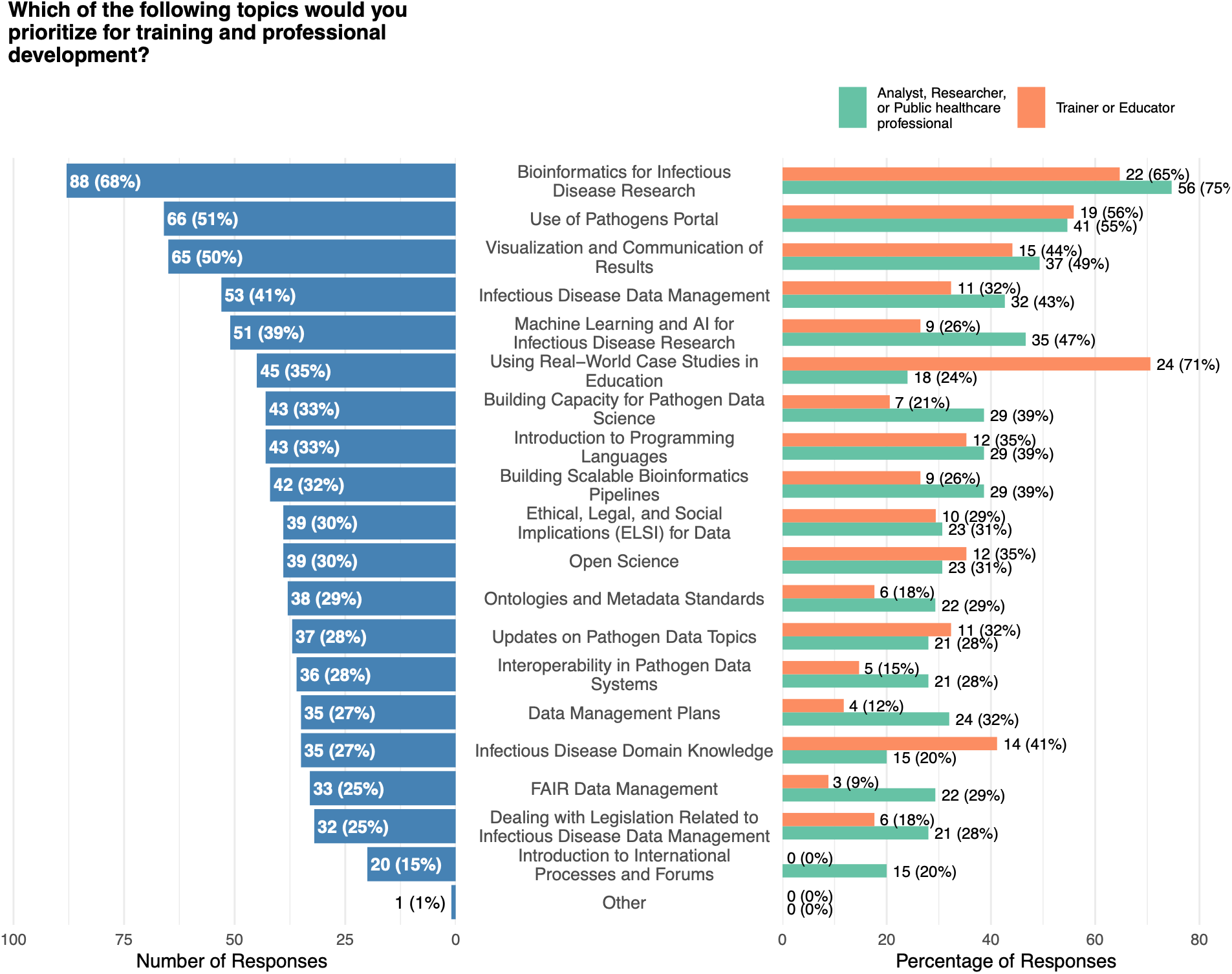
Results of the multiple select question on training prioritization, summarized over all responses (left) and split by the most frequently indicated professional roles (‘Analyst, Researcher or Public healthcare professional’ and ‘Trainer or Educator’; right).

Preferences for how training should be delivered were also captured (Fig 4). Virtual short courses were the most popular format, with 68 % of respondents indicating a preference for this mode of instruction.

**Fig 4.**
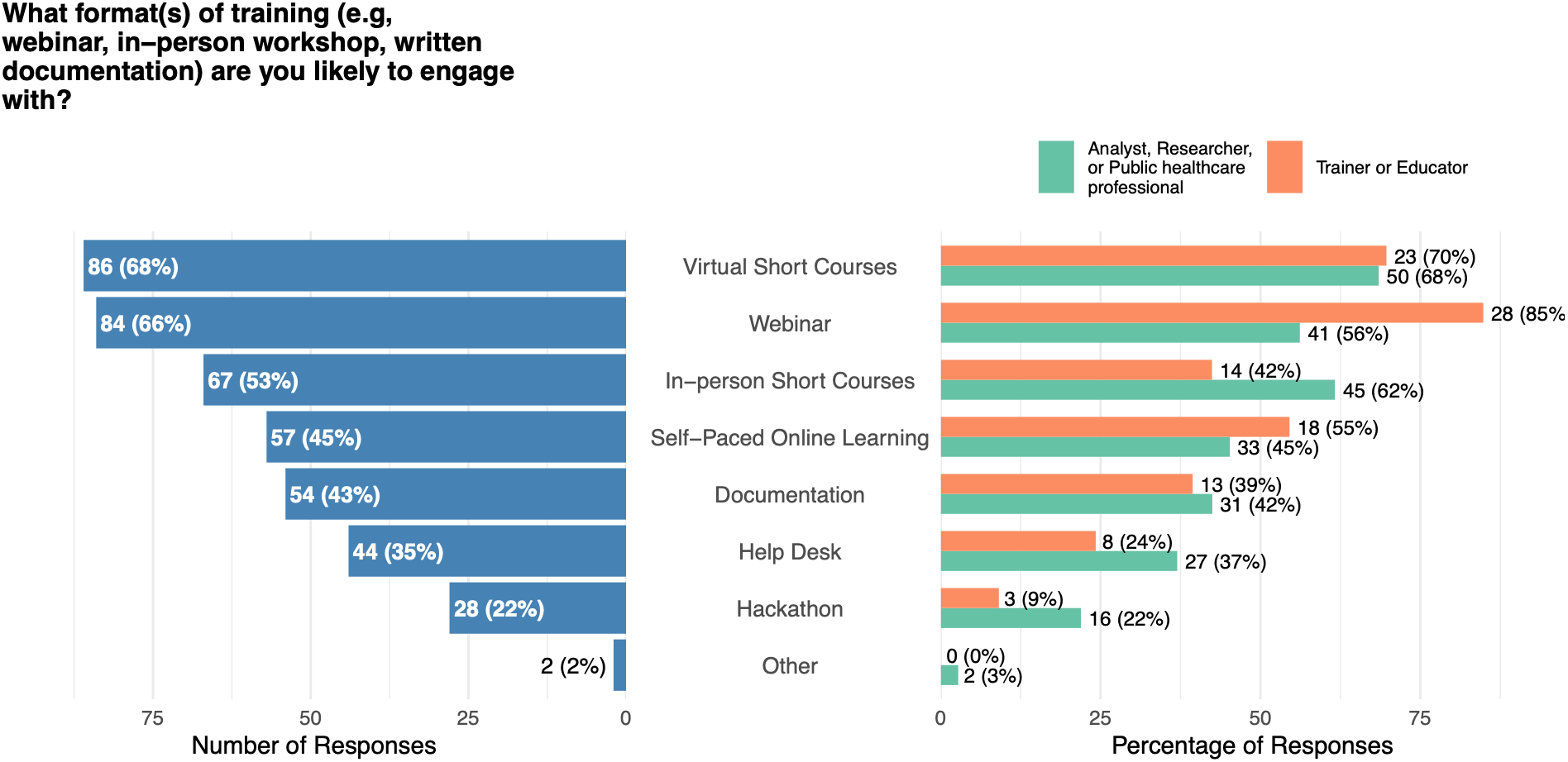
Results of the multiple select question on training format preference, summarized over all responses (left) and split by the most frequently indicated professional roles (‘Analyst, Researcher or Public healthcare professional’ and ‘Trainer or Educator’; right).

Webinars were close behind (66 %), followed by in-person short courses (53 %). Self-paced online learning materials attracted 45 % of the sample, while written documentation such as manuals or guides was preferred by 43 %. These preferences were broadly consistent across both researchers and educators, although educators showed a higher interest toward webinars and a lower interest toward in-person courses.

The survey also asked participants to rank the importance of various resources provided by the PDN (Fig 5). The Pathogen Portal was identified as the most valuable resource, with 72 % of respondents indicating that it was essential to their work or that of their peers. Metadata standards were considered important by 43 % of participants, while validated analytical workflows and access to expert advice each received endorsement from 37 % of the sample. In-person workshops aimed at undergraduate students were valued by 35 % of respondents.

**Fig 5.**
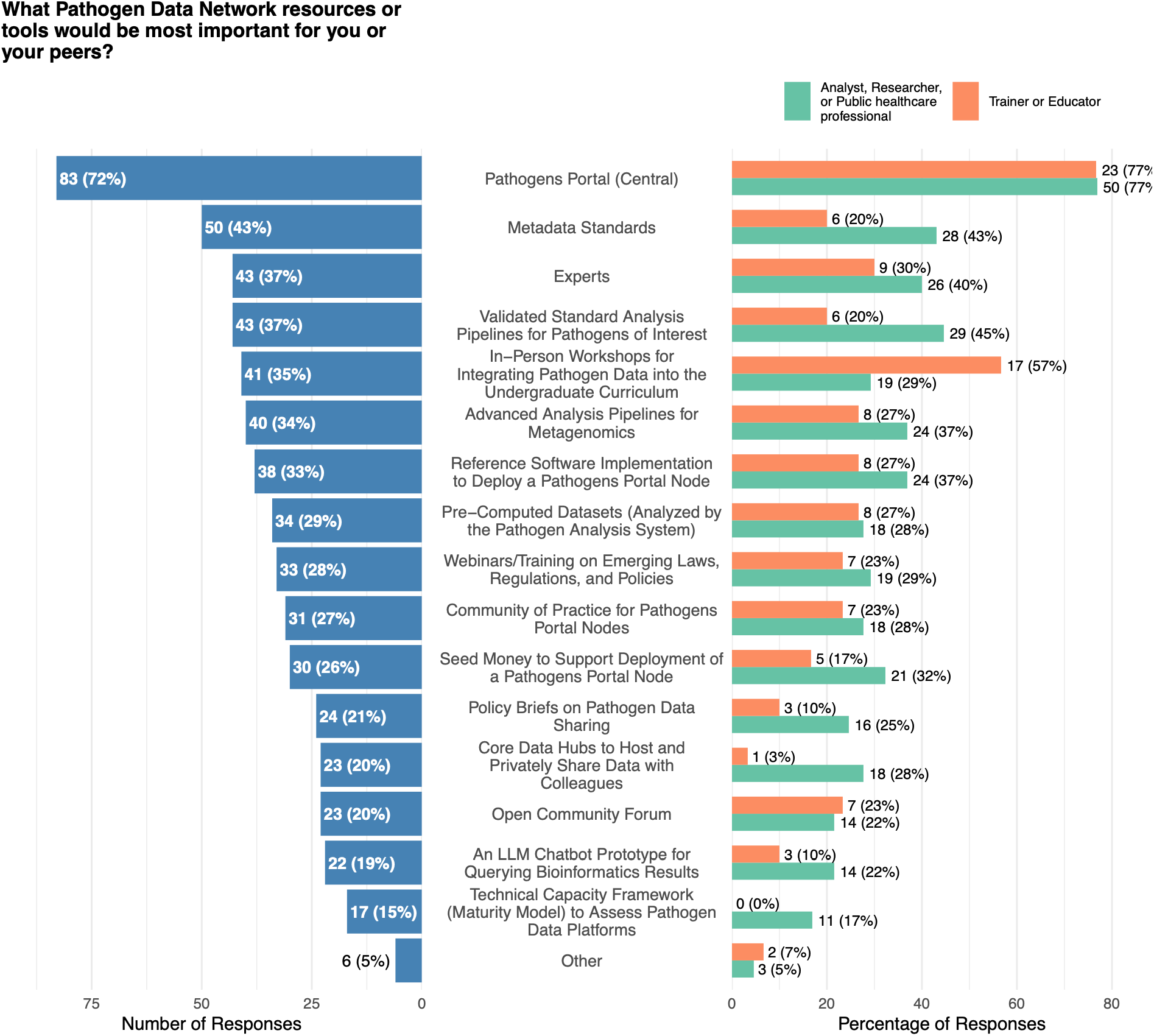
Results of the multiple select question on the importance of PDN resources or tools, summarized over all responses (left) and split by the most frequently indicated professional roles (‘Analyst, Researcher or Public healthcare professional’ and ‘Trainer or Educator’; right).

## Discussion

This survey provides a structured assessment of how professionals working with pathogen data perceive their operational environment, resource constraints, and workforce development needs. Although the findings are descriptive, several patterns emerge consistently enough to inform planning for data infrastructure and training initiatives within the infectious-disease research ecosystem.

The predominance of respondents identifying limited funding (74%), difficulties with data aggregation (68%), and shortages of skilled personnel (52%) as significant barriers points to an enduring imbalance between the complexity of pathogen-data tasks and the resources available to carry them out. [6,13] Similar observations have been reported in bioinformatics and infectious-disease research more broadly, where the rapid expansion of genomic and epidemiological data has outpaced investments in infrastructure and workforce development [7,12]. Notably, relatively few respondents cited technological limitations or concerns about data misuse as primary barriers. This pattern suggests that the most immediate constraints on FAIR pathogen data are organizational and structural rather than technological—sustaining skilled personnel, coordinating across institutions, and maintaining stable financial support.

Training priorities reinforce this interpretation. Two-thirds of respondents identified bioinformatics for infectious-disease research as their highest training priority, followed by training on the Pathogens Portal (51%) and on data visualization and communication (50%). Similar to previous studies [13–15], these preferences delineate a workforce that is aware of computational needs but still developing competence in integrating, interpreting, and presenting data. Also, machine learning and AI for infectious disease research was frequently mentioned as training priority (39%). This is an area that is strongly gaining importance in all aspects of society, and therefore important to address.

While in the past survey outcomes pointed towards preferences for short in-person courses [13], we saw more emphasis on online delivery modes—68% selecting virtual short courses and 66% selecting webinars—which reinforces the value of flexible, distributed instruction formats, though such preferences may also reflect current constraints on time, funding, or institutional support rather than an intrinsic pedagogical choice.

Differences between researchers and educators, while expected, are instructive. Researchers emphasized technical and procedural aspects of data stewardship, particularly FAIR-data management and planning, whereas educators focused on conceptual understanding and pedagogical framing. The contrast suggests that pathogen data occupy distinct functional roles in research and teaching: one as an analytical substrate, the other as a medium for instruction. Recognizing this distinction will be important for any coordinated training initiative.

Responses regarding the Pathogen Data Network’s resources reinforce the same structure of need. The integrated pathogen data and tools portal (Pathogens Portal) was identified by 72% of participants as the most important PDN output, followed by metadata standards, validated workflows, and access to experts. These results imply that the community continues to view centralization, standardization, and mentorship as the most tangible support for effective data use. While respondents did not comment directly on governance, the relatively lower concern with data misuse may reflect evolving trust in established data sharing frameworks, or simply limited engagement with governance frameworks.

Several limitations should be considered when interpreting these findings. The survey was conducted in English and distributed primarily through established research networks, and responses resulted from convenience sampling. This likely introduced selection bias toward individuals already engaged with pathogen-data initiatives. The instrument was intentionally brief, consisting of ten questions, and was designed to capture broad patterns rather than support inferential statistical analysis. Consequently, the results should be interpreted as a descriptive snapshot rather than a statistically representative sample of the entire pathogen-data community. Future work would benefit from longitudinal assessments, multilingual deployment, and integration with empirical measures of data-sharing practices to determine whether perceived barriers correspond to observed behavior.

Despite the above-mentioned constraints, the results provide an empirical baseline for evaluating the operational state of pathogen-data research and education. The reported convergence on funding, aggregation, and workforce shortages suggests that the immediate impediments are structural and systemic rather than conceptual. Addressing them will require sustained investment in training, coordination, and institutional continuity, alongside developing adequate technical tools. The patterns observed here should therefore inform—not dictate—the priorities of any organization seeking to improve the interoperability and reusability of pathogen data, including the Pathogen Data Network. The survey questions themselves can also help establish a foundation for future community assessments. For example, through its “Open Community Forum,” the PDN will be supporting broad community feedback-gathering exercises which can build on the findings of this survey work.

## Conclusions

The outcomes of the PDN stakeholder survey suggest there is a major need for support in retrieving, analyzing, federating and interpreting pathogen data from the academic community. Providing this support is not only of utmost importance to tackle current and future infectious-disease outbreaks, but lso to answer more fundamental questions in pathogen research.

## Acknowledgements

This project has been supported in whole or in part by the National Institute of Allergy and Infectious Diseases of the National Institutes of Health under Award Number U24AI183840.

## Notes

https://doi.org/10.5281/zenodo.19494321

https://github.com/sib-swiss/pdn_survey_analysis

### Competing Interest Statement

The authors have declared no competing interest.

### Summary of Updates

After review, we updated the manuscript. The title, abstract and introduction now state that this was an open, cross-sectional convenience survey with an unknown eligible population and no calculable response rate. The target community is defined as researchers, educators, healthcare professionals and policymakers. The three-point barrier scale is no longer called a Likert scale, and 'diagnostic' is now 'descriptive'. The methods now say the survey was reported in line with the CROSS checklist, and a CROSS checklist has been filled in against the revised text.

